# Granular retrosplenial cortex layer 2/3 generates high-frequency oscillations coupled with hippocampal theta and gamma in online states or sharp-wave ripples in offline states

**DOI:** 10.1101/2023.07.10.547981

**Authors:** Kaiser C. Arndt, Earl T. Gilbert, Lianne M. F. Klaver, Jongwoon Kim, Chelsea M. Buhler, Julia C. Basso, Sam McKenzie, Daniel Fine English

## Abstract

Neuronal oscillations support information transfer by temporally aligning the activity of anatomically distributed ‘writer’ and ‘reader’ cell assemblies. High-frequency oscillations (HFOs) such as hippocampal CA1 sharp-wave ripples (SWRs; 100-250 Hz) are sufficiently fast to initiate synaptic plasticity between assemblies and are required for memory consolidation. HFOs are observed in parietal and midline cortices including granular retrosplenial cortex (gRSC). In ‘offline’ brain states (e.g. quiet wakefulness) gRSC HFOs co-occur with CA1 SWRs, while in ‘online’ states (e.g. ambulation) HFOs persist with the emergence of theta oscillations. The mechanisms of gRSC HFO oscillations, specifically whether the gRSC can intrinsically generate HFOs, and which layers support HFOs across states, remain unclear. We addressed these issues in behaving mice using optogenetic excitation in individual layers of the gRSC and high density silicon-probe recordings across gRSC layers and hippocampus CA1. Optogenetically induced HFOs (iHFOs) could be elicited by depolarizing excitatory neurons with 100 ms half-sine wave pulses in layer 2/3 (L2/3) or layer 5 (L5) though L5 iHFOs were of lower power than in L2/3. Critically, spontaneous HFOs were only observed in L2/3 and never in L5. Intra-laminar monosynaptic connectivity between excitatory and inhibitory neurons was similar across layers, suggesting other factors restrict HFOs to L2/3. To compare HFOs in online versus offline states we analyzed, separately, HFOs that did or did not co-occur with CA1 SWRs. Using current-source density analysis we found uniform synaptic inputs to L2/3 during all gRSC HFOs, suggesting layer-specific inputs may dictate the localization of HFOs to L2/3. HFOs occurring without SWRs were aligned with the descending phase of both gRSC and CA1 theta oscillations and were coherent with CA1 high frequency gamma oscillations (50-80 Hz). These results demonstrate that gRSC can internally generate HFOs without rhythmic inputs and that HFOs occur exclusively in L2/3, coupled to distinct hippocampal oscillations in online versus offline states.

## Introduction

The theory of systems consolidation of memory posits that internal representations of experience in hippocampal (HPC) CA1 steer reorganization of distributed neocortical cell assemblies to create a long-term memory of events^1–3^. During online states (e.g. ambulation through space; associated with high cholinergic tone) local field potentials, spiking of neuronal populations, and membrane potentials of individual neurons in both hippocampal and downstream cortical networks are dominated by theta and gamma activity^4–8^. In contrast, offline states (e.g. quiet wakefulness; associated with diminished cholinergic signaling) are highlighted by CA1 sharp-wave ripples (SWRs)^9–11^, which are also reflected in the local field potential (LFP), neuronal spiking, and single neuron membrane potentials^5–8,12,13^. CA1 theta oscillations organize place cells into assemblies during experience, and by modifying synaptic connectivity are believed to enable subsequent SWRs to function as a “teaching signal” to induce plasticity in downstream cortical regions to consolidate the memory of the experience^2,3,14^. Cortical high-frequency oscillations (HFO; 100-250 Hz; most prevalent in midline and parietal cortices) likely support the processing of coincident hippocampal SWRs.

One region that exhibits HFOs is the granular retrosplenial cortex (gRSC). Changes in gRSC oscillatory activity are observed in several neurological and neuropsychiatric disorders^15–17^. Functionally, the gRSC supports spatial navigation as well as memory, and contains place, head direction, boundary vector, and border cells^18–23^. Mechanisms of HFO generation including laminar organization, correlation with specific brain states, and coupling to hippocampal oscillations remain unclear.

We addressed these gaps using anatomically registered laminar local field potential and single unit recordings combined with focal optogenetics in behaving mice. We found that activation of excitatory neurons in either layer 2/3 (L2/3) or layer 5 (L5) of gRSC induced HFO-like activity (iHFOs), though spontaneous HFOs and associated synaptic currents were only observed in L2/3. Because monosynaptic connectivity was found to be similar across layers, layer specific inputs appear critical in supporting HFOs. We found that in online states HFOs preferentially occurred at the descending phase of theta oscillations and were coherent with CA1 gamma, while in offline states HFOs characteristically occurred with sharp-wave ripples. These findings suggest that the ability for gRSC to intrinsically generate HFOs may enable coupling with different HPC oscillations during different behaviors.

## Results

### HFOs are localized to gRSC Layer 2/3 across states

To investigate intra- and inter-regional communication in gRSC and HPC circuits we obtained LFP and single unit recordings in awake head-fixed mice, navigating a 1-D visual virtual environment by running on top of a wheel. One high density linear silicon probe (20 μm pitch spanning 1200 μm) was implanted across the hemispheres beneath the central sinus spanning the layers of the gRSC, with a second probe inserted through ipsilateral HPC CA1 (**Fig. 1A**). Probe locations were identified online using electrophysiological landmarks and post-hoc using Di-I staining and probe tracks in the tissue (**Fig. 1B**). To examine layer specific oscillatory power, we computed power spectral densities (PSDs) across anatomical space for entire recording sessions (i.e. including both ambulation/navigation through virtual space and quiet wakefulness; **Fig. 1C**.) HFO (100-250 Hz) power was highest in L2/3, weakly extending into L5. To anatomically localize current sources and sinks at the time of HFOs we detected events in L2/3 and computed HFO-triggered CSDs which revealed rhythmic current sources and sinks localized to L2/3 (**Fig 1D**). Detecting HFOs from L5 LFP gave the similar results (**Supplementary Fig. 1**), confirming our finding is not due to event detection location. Furthermore, events detected in each layer coincided in time and were both associated with higher HFO power in L2/3, consistent with the interpretation that HFOs in the L5 LFP are not independent from L2/3 events and are likely due to volume conduction.

**Fig. 1:**
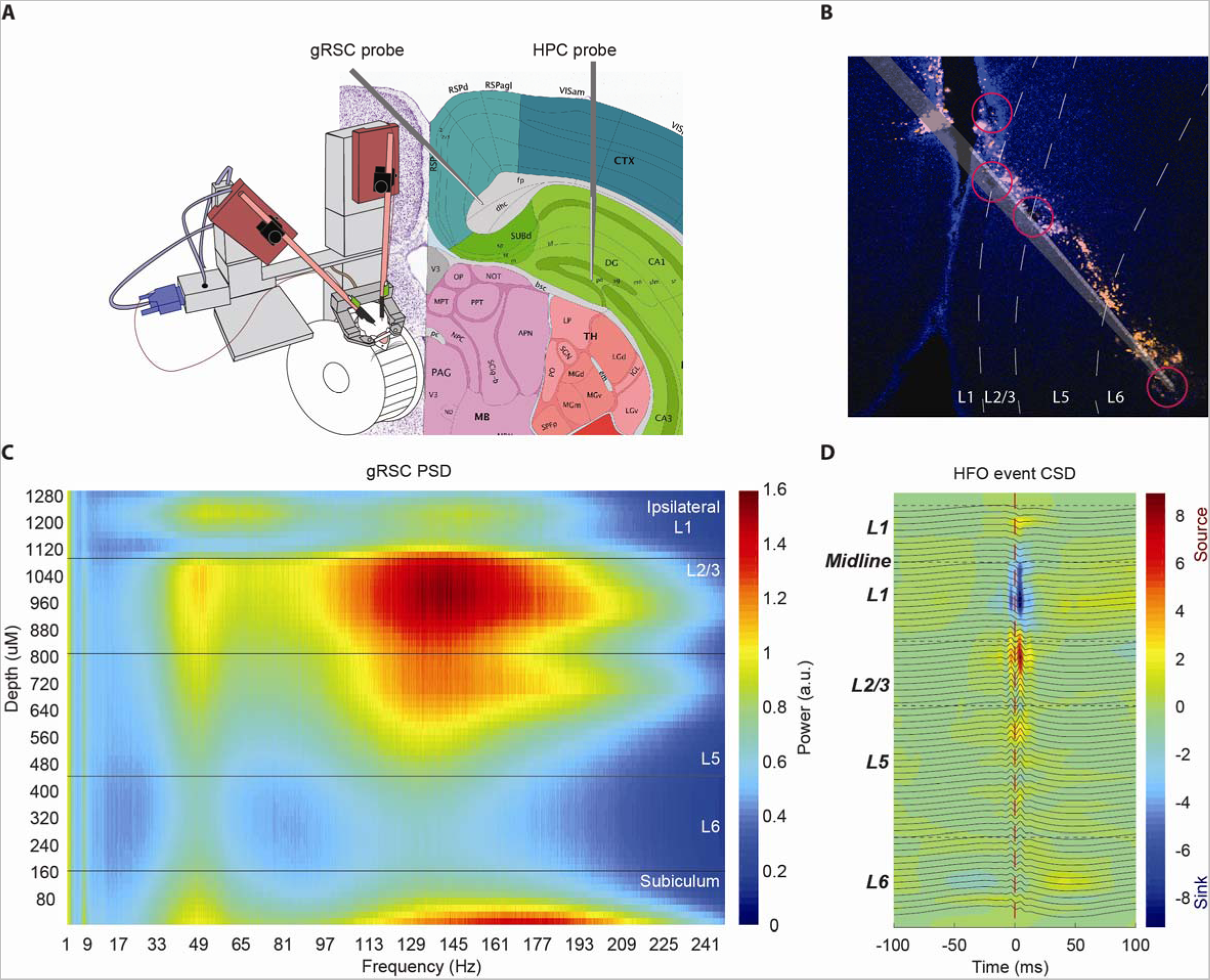
High frequency oscillation activity is highest in layer 2/3 across all brain states: **A)** Graphic of acute head-fixed recording and multi-probe implant locations in the gRSC and HPC CA1. **B)** Photomicrograph of the gRSC of a single session implant track (orange, Di-I) and electrolytic lesions (red circles) delivered through the silicon probe to confirm layer locations. **C)** Representative power spectral density plot across all layers of the gRSC. **D)** Representative event triggered current source density plot for all detected HFOs from the L2/3 channel, time zero is the peak of the HFO events.

### Ultra-Focal optogenetic excitation generates higher power HFOs in L2/3 as compared to L5

To gain insight into the possible role of each layer in generating HFOs, we tested the ability of each layer to generate HFOs when optogenetically activated (similar manipulations have been performed in the HPC CA1)^12^. To accomplish this we used ultra-focal, layer specific, optogenetic stimulation of excitatory neurons with 4-shank 32-channel μ LED probes^24^, with shanks in L2/3 and L5 (**Fig. 2A**). Induced HFOs (iHFOs) were observed in both L2/3 and L5 (p < 0.0001) in response to optical stimulation (100 ms half-sine wave pulses applied semi-randomly at 2-3 second intervals). Importantly, responses were restricted to the stimulated layer, confirming our ability to stimulate each layer independently (**Fig. 2B**). L2/3 iHFOs were of significantly higher power than L5 iHFOs (Wilcoxon rank sum test, p < 0.0001), consistent with results from our PSD analysis (**Fig. 2C**). These results suggest that while spontaneous HFOs are restricted to L2/3, non-rhythmic activation of either L2/3 or L5 local circuits generates intralaminar iHFOs.

**Fig. 2:**
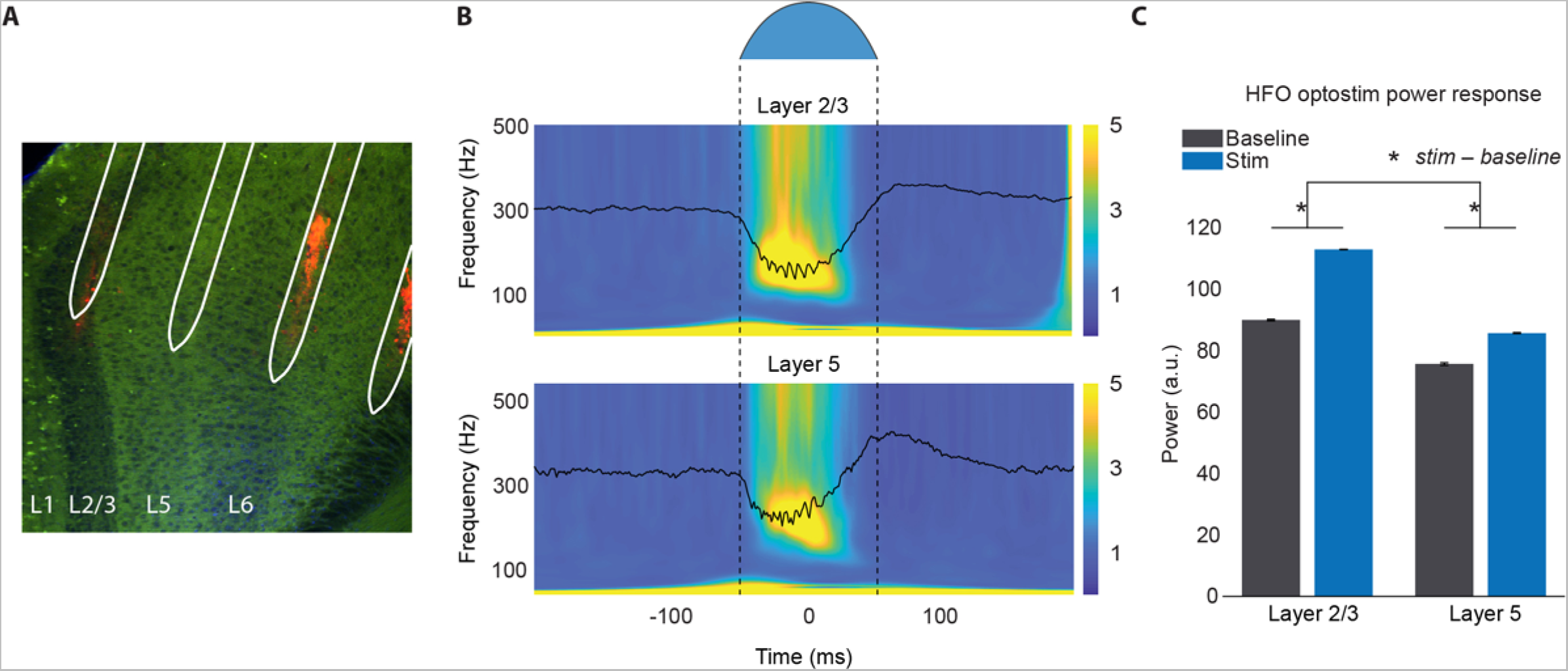
Both gRSC L2/3 and L5 are capable of generating HFOs: **A)** Photomicrograph of the gRSC of CaMKII-Cre::Ai32 mice. White overlay of Neurolight Technologies’ µLED probe. Channelrhodopsin (green) expressed in excitatory cells. Di-I (orange) used to mark the tracks of the probe shanks. **B)** *Top*: shape of 100 ms light stimulation pulse delivered to each layer of the gRSC. Light intensity was determined empirically in L2/3 and set to a level that reliably evoked HFOs. *Middle*: Representative L2/3 optical stimulation event-triggered PSDs (150 pulses to each layer) z-scored and normalized to the 500 ms before analysis window shown. Overlaid traces are the event-triggered average LFP. Time 0 is the peak of the stimulation pulse. Bottom: Same as middle, but for L5. **C)** Averaged power response in the 100-250 Hz range during baseline and middle of the stimulation. Both L2/3 and L5 reliably generated HFOs during stimulations (baseline compared to stim; Wilcoxon signed rank test, p < 0.0001). Difference of stim to baseline in HFO power is significantly higher in L2/3 compared to L5 (Wilcoxon rank sum test, p < 0.0001) (3 animals, 150 light pulses to each layer in each animal).

### Characteristics of HFOs across states

gRSC HFOs are coupled with CA1 sharp wave-ripples, and events of similar frequency are entrained to theta oscillations, particularly in REM sleep. ^25–30^. To compare the physiology of these events we separately analyzed HFOs in online and offline states (distinguished by characteristic CA1 oscillations) HFOs occurring within +/− 50 ms of a CA1 SWR were designated as with-SWR events (HIR) and those outside this window as outside-SWR events (HOR) (**Fig. 3A, B**). HORs were significantly longer than HIRs (n=7720 HIRs, 71965 HORs; p-value < 0.0001), while HIRs had significantly higher amplitude, frequency, and power (p-value < 0.0001) (**Fig. 3C, D**). We next quantified the spiking activity of putative excitatory neurons (ENs) and interneurons (INs) using CellExplorer^31^ (**Supplementary Fig. 2**). Neurons were assigned to layers by estimating their position using the recording site with maximum spike amplitude. L5 ENs and INs both increased their firing rate more during HORs than HIRs (ENs, p < 0.01; INs, p < 0.01), and L1 interneurons increased their firing rate more during HIRs than HORs (p = 0.045). In contrast the gain in L2/3 firing rate was consistent across HIRs and HORs (**Fig. 3E**). These results suggest L2/3 activity is critical for all HFOs.

**Fig. 3:**
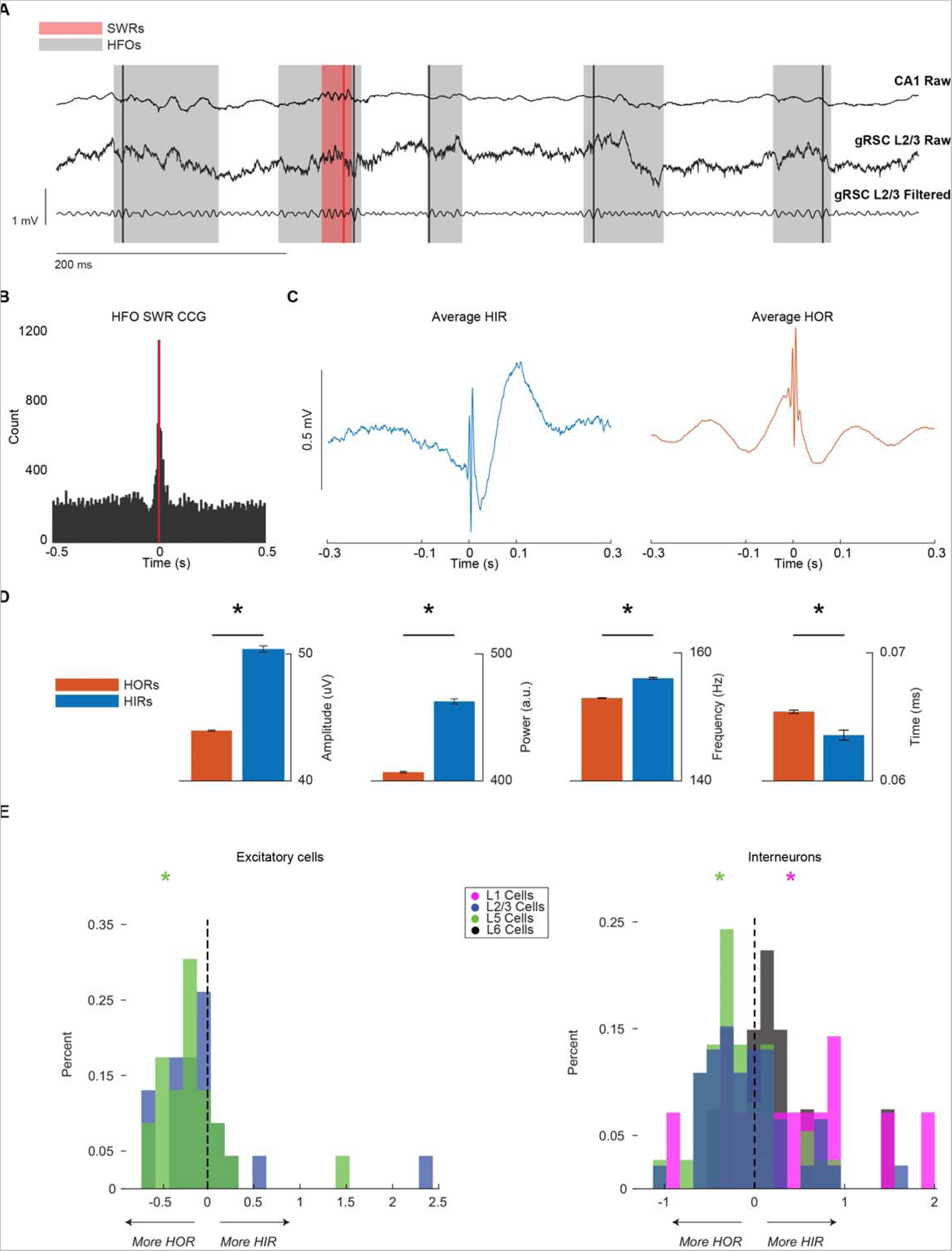
High frequency oscillations are identified during and outside of SWRs: **A)** Example traces and detected SWRs (red) and HFOs (grey) in CA1 and gRSC respectively. Lines mark the peak of the detected event; shaded areas mark the duration of the detected event. **B)** SWR-HFO cross correlogram. Red line marks the peak of detected SWRs. **C)** *Left*; single session representative average HFOs in ripples (HIRs) LFP trace (513 events). *Right*; same as left but for HFOs out of ripples (HORs) (14986 events). **D)** Comparison of amplitude, power, frequency, and duration (left to right) between HIRs and HORs across 4 recordings from 4 animals (7720 HIRs; 71965 HORs). Asterisks represent p-value < 0.001. An unpaired T-test was used for HFO frequency distributions, all others used Wilcoxon Rank Sum test. **E)** Gain in firing rate for putative excitatory cells (left) and interneurons (right) during HIRs and HORs. X axis is the difference between the gain in firing during HIRs and HORs (HIR gain – HOR gain). Colored asterisks correspond to the layer they were recorded in and designate whether those units fired more during either event type. Layers without asterisks did not have cells that fired more during either HIRs or HORs.

### L2/3 and L5 have similar intra-laminar connectivity

To determine if connectivity patterns could explain why HFOs are localized to L2/3, we quantified putative excitatory and inhibitory connections by examining auto- and cross-correlograms for spike trains of gRSC neurons (**Fig 4A**), both within and between layers (**Fig. 4B**). 65.6% of monosynaptic connections were observed to be intra-laminar. Since both L2/3 and L5 are capable of generating local HFOs and local monosynaptic connections are thought to support HFO generation, we quantified the percent of ENs and INs with a connection to another cell within the presynaptic cell’s layer; termed E-E, E-I, I-I, and I-E. Connection probabilities were not different across L2/3, L5, and L6 (ANOVA analysis, E-I p = 0.90; I-I p = 0.52; I-E p = 0.47). L1 was excluded from this analysis as no intralaminar monosynaptic connections were identified. E-E connections were also excluded as only 3 putative connections were identified. The similar excitatory and inhibitory connectivity likely facilitates iHFO generation in each layer (**Fig. 2**). However, since endogenous HFOs don’t occur in L5 our findings here also suggest that inputs to each layer are distinct. i.e., L5 does not receive the correct source or level of input to generate endogenous HFOs.

**Fig. 4:**
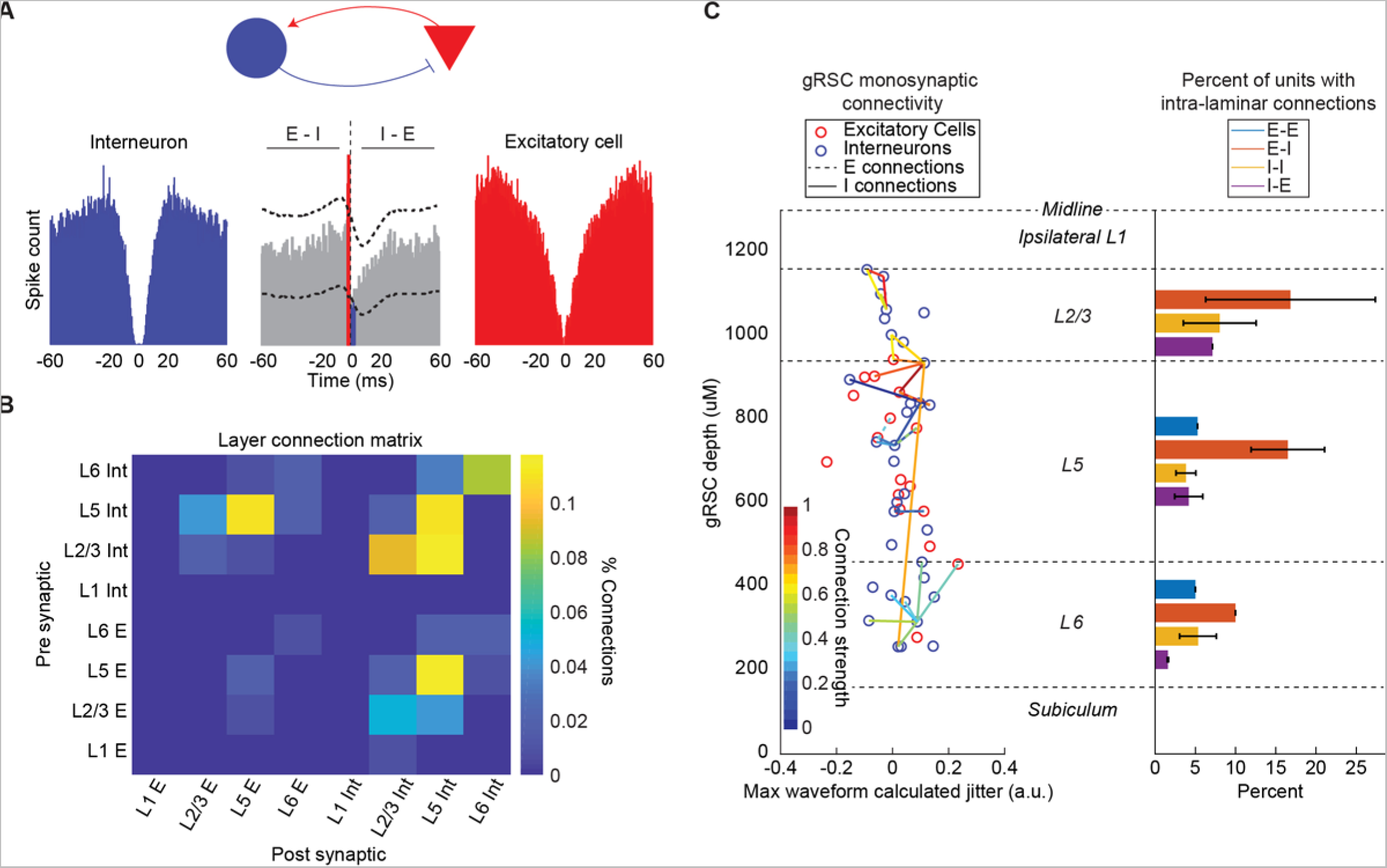
Inter and intra-laminar monosynaptic connections in the gRSC: **A)** Example monosynaptic excitatory and inhibitory connection between a putative excitatory and inhibitory cell. *Middle;* Cross correlogram between the interneuron and excitatory cell. The peak before zero shows excitatory drive of the interneuron (I) (red bars) and the trough after zero shows the inhibition of he excitatory cell (E). Horizontal dotted lines are the 95% confidence interval of the average cross correlation at each time point. **B)** Connectivity matrix for the percent of monosynaptic pairs between Es and Is across the layers of the gRSC. **C)** *Left;* Representative monosynaptic connectivity map showing the putative locations of each unit and their monosynaptic connections across the layers of the gRSC. *Right;* Percent of each cell type with putative intra-laminar connections in each layer. Bars not shown in each layer had no detected monosynaptic connections of that type.

### HFOs in both online and offline states are associated with CSD sinks in L2/3

HIR and HOR event-triggered PSDs revealed differences in associated rhythmic network activity. HIRs were associated with a negative LFP deflection indicative of excitatory inputs (**Fig. 5A**), along with high delta (1-4 Hz) power and lack of movement (characteristics of offline states). HORs occurred when gRSC and CA1 exhibited LFP theta oscillations, preferentially occurring at the descending phase of the oscillation (**Fig. 5B**). This rhythmic theta activity is clearly present in both the HOR-triggered LFP (**Fig. 3C**) and PSD (**Fig. 5A**). While the rate of HORs was similar for periods of ambulation vs stillness, nearly all HIRs occurred when the animals were still (**Fig. 5C**). Additionally, animal velocity and theta power during HORs in both gRSC and HPC was significantly higher than during HIRs (**Supplementary Fig. 3**). CSDs revealed rhythmic current sources and sinks located on the border between ipsilateral L2/3 and L1 for both HIRs and HORs. (**Fig. 5D**). To determine if these results were biased by detection of HFOs in L2/3 and not L5 (a smaller amplitude HFO is seen in L5 at the same time) we detected HFOs from L5. We found current sources and sinks were also located flanking the L1-L2/3 border (**Supplementary Fig. 1**), suggesting apparent L5 HFOs reflect volume conduction from L2/3. These data, combined with the results from ultra-focal stimulation of each layer (**Fig. 2B**) and monosynaptic connectivity analyses (**Fig. 4**) support the conclusion that the ability for a layer to generate spontaneous HFOs is controlled by their relative inputs.

**Fig. 5:**
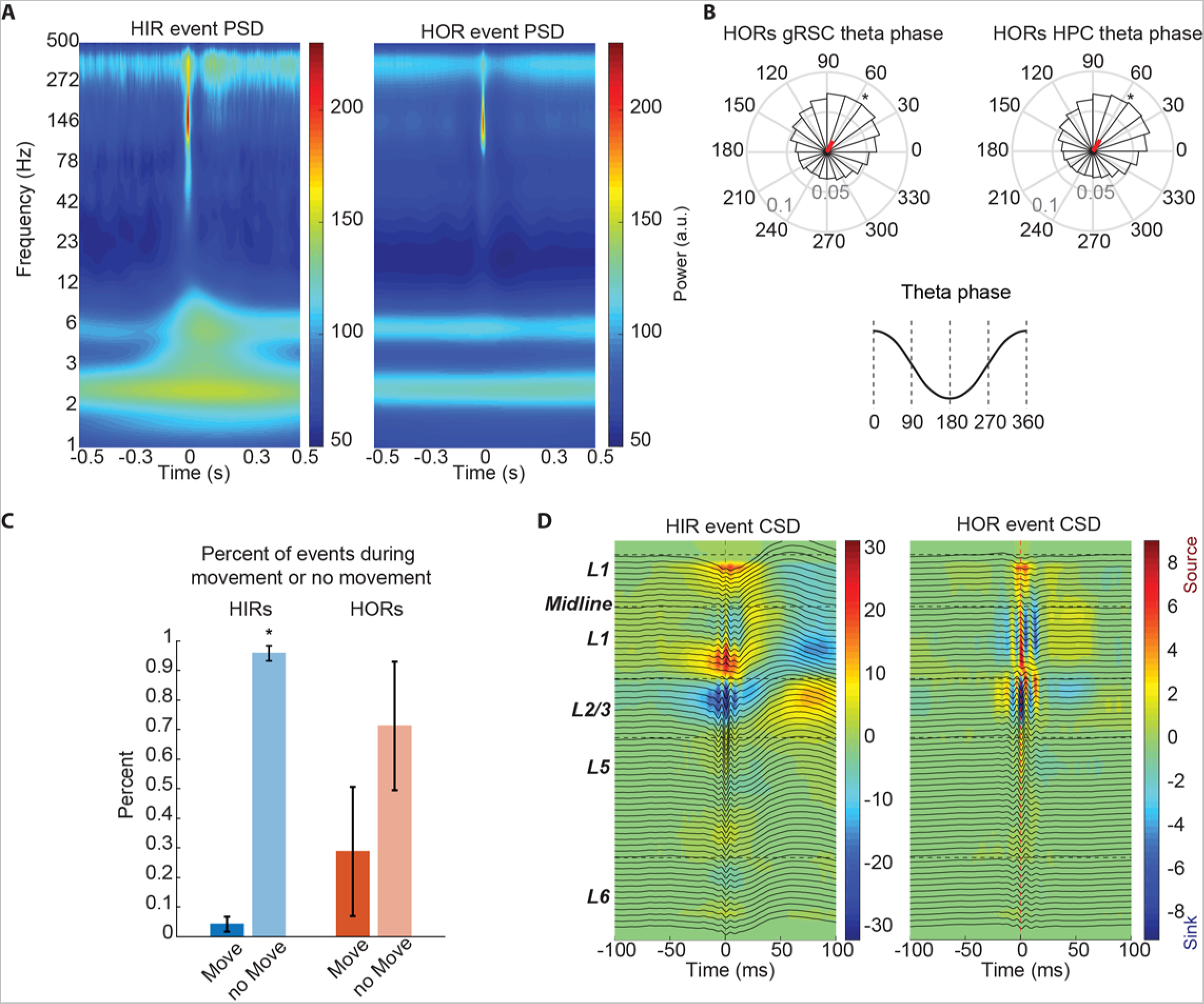
HFO activity is on the boarder of L1-L2/3 and HORs occur on the descending phase of local and HPC theta: **A)** *Left*; Representative HIR (1128 events) event-triggered power spectral density (PSD) in the gRSC. The peak of each event is centered on 0. *Right*; Same as left but for HORs (30396 events) from the same session. **B)** Polar plots of HOR (4 animals, 87369 events) preference to local gRSC theta (left) and HPC theta (right). Both significantly occur on the descending phase of theta (Rayleigh’s test p < 0.0001). **C)** Percent of HIRs (blue) and HORs (orange) that occur when the animal is stationary or moving (Wilcoxon rank sum test p < 0.05). **D)** *Left*; Representative HIR event-triggered current source density (CSD) centered the same and using the same events as in A. *Right*; Same as left but for HORs from the same session.

### HPC high gamma frequency is rhythmic and coherent with HORs

To determine the oscillatory activity of CA1 circuits during HIRs and HORs we computed HIR and HOR triggered PSDs and CSDs from the CA1 pyramidal layer LFP (**Fig. 6A, B**). As expected, we observed increased SWR-frequency power in HIRs along with characteristic current sources and sinks in Str. Pyr and Str. Rad, respectively (**Fig. 6A, B *left***). In HORs, pyramidal layer LFP was characterized by a persistent increase in theta power and theta-nested gamma. Gamma activity was significantly stronger in the high gamma range (~50-80Hz, N=87369 HORs) (p < 0.0001; **Fig. 6C**). Consistent with previous reports CA1 theta activity was associated with current sources and sinks in the pyramidal layer and Str. L.M., respectively (**Fig. 6B**)^32,33^. Further analysis revealed significant power-power coupling between the gRSC HFO frequencies and CA1 high gamma (**Fig. 6D**). This spectral coupling suggests gRSC HORs and HPC high gamma could be frequencies used for communication and experience-dependent information transfer during online states^33,34^.

**Fig. 6:**
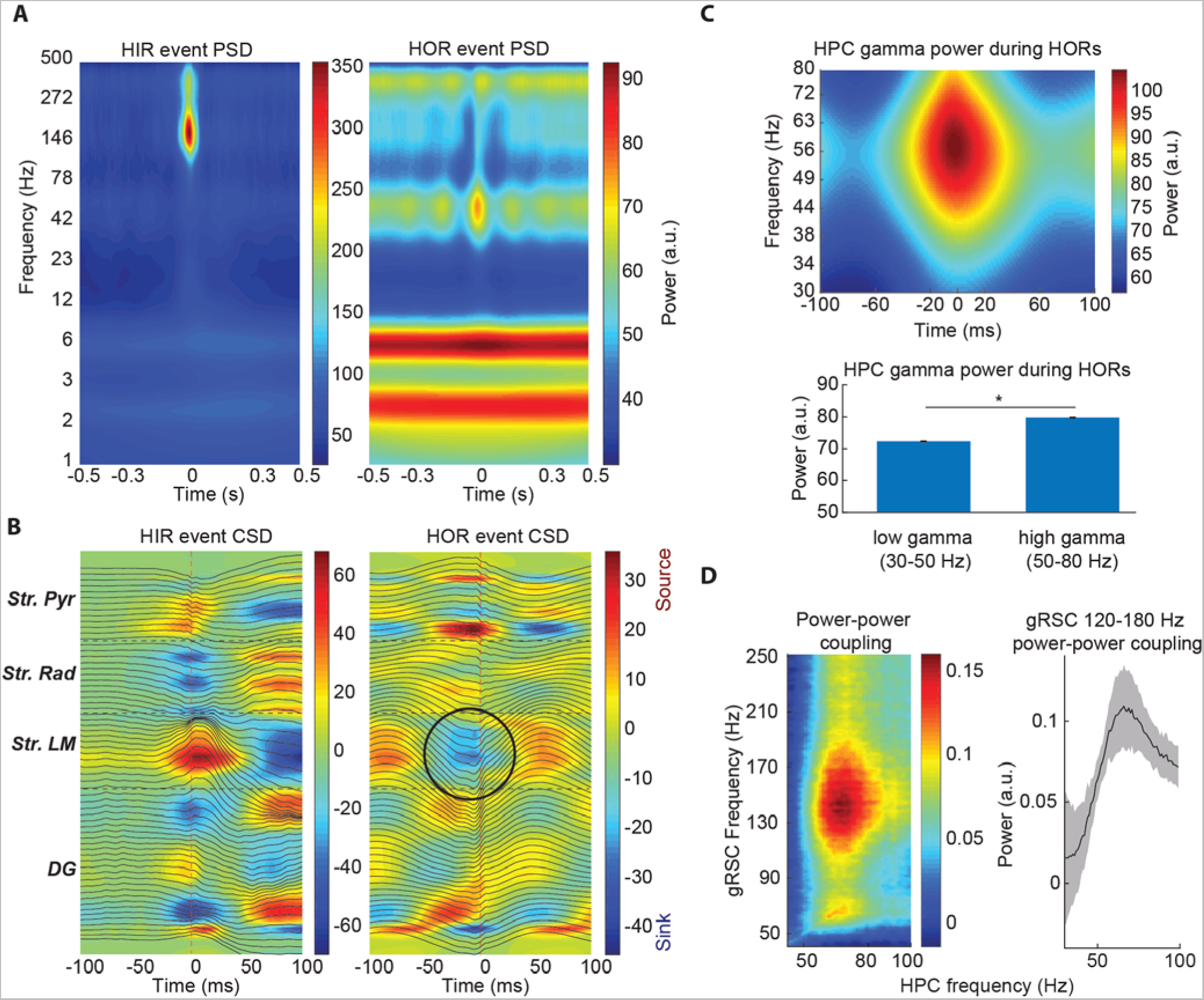
Spectral coupling dynamics of HPC gamma and SWRs with gRSC HFOs: **A)** *Left*; Representative HIR (1128 events) event-triggered PSD in the HPC channel used to detect SWRs. 0 is centered on the peak of the HIR. *Right*; Same as left but for HORs (30396 events). **B)** *Left*; Representation HIR event-triggered CSD centered the same and using the same events as in A. *Right*; Same as left but for HORs. Black circle marks current sink in Stratum L.M. **C)** *Top*; Averaged gamma band PSD centered on HOR peaks (4 animals). *Bottom*; Difference in power of low and high gamma bands during HORs (p < 0.0001) (87369 HORs). **D)** *Top*; Representative spectral coupling between HPC channel used to detect SWRs and gRSC channel used to detect HFOs. *Bottom*; Average spectral coupling of the gRSC 120-180 Hz to HPC frequencies (4 animals).

## Discussion

In rodents and humans, high-frequency cortical oscillations are prevalent across online and offline states in cortical regions receiving excitatory drive from the hippocampal formation^35–38^, which suggests flexible coupling of these events to different hippocampal oscillations^39^. In offline states HFOs are temporally aligned with SWRs, while in online states HFOs switch to being aligned with CA1 theta/gamma oscillations^26,27,40^ (**Fig 6**). Here we investigated mechanisms of HFO generation across brain states, including layer specific gRSC physiology and its coupling to CA1 activity.

### gRSC HFOs are generated exclusively in L2/3

HFOs have been reported to occur throughout broadly defined anatomical locations including “superficial” or “deep” gRSC^26,27,29,40^, making their cellular basis unclear. Here we mapped the laminar electro-anatomy of the gRSC and found that HFOs occur exclusively in L2/3. This is supported by our current source density (CSD) analysis showing that during all events (HIRs and HORs), the current sources and sinks flank the L1-L2/3 border. HFOs are thus driven by currents arriving in L2/3. Recently gRSC ‘splines’, with similar frequencies to HFOs, have been described in superficial gRSC, aligned with theta oscillations. We believe that ‘splines’ and HFOs are one and the same, and since we observed these events to be physiologically consistent in both theta and SWRs, we suggest that HFOs are a general mechanism by which gRSC circuits process hippocampal inputs (CITE).

### Synaptic inputs rather than intralaminar connectivity restrict spontaneous HFOs to L2/3

Intralaminar connectivity, in particular E-I connections known to be important for rhythmogenesis, was not found to be different between layers, and broadband stimulation of either L2/3 or L5 resulted in iHFOs. The lack of spontaneous HFOs in L5 is likely due to differential inputs to each layer during theta and SWRs, as demonstrated by current sinks and sources being restricted to L2/3 in both theta (HORs) and SWRs (HIR). We found that excitatory and inhibitory L5 units both increased their firing rate more during HORs than HIRs, potentially facilitated by long-range inhibitory projection neurons from CA1 connecting to gRSC L1 projecting L5 pyramidal neurons^41^. Additionally, L5 packet activity during cortical synchronous states co-terminate with SWRs, also potentially through these inhibitory projection neurons^40^. Given these previous works paired with our findings, we conclude L5 units present a higher firing rate during HORs due to being more suppressed during HIRs^27^.

We also report L1 interneurons exhibit greater firing rate during HIRs than HORs. As previously reported in Nitzan et al., 2020, VGtlut2^+^ bursting neurons in the subiculum neurons project to superficial gRSC, notably to both L1 and L2/3. Since excitatory neurons are not present in L1^42,43^, this increase in firing rate during HIRs is likely driven by inputs from the subiculum.

### Coupling of HFOs to CA1 oscillations

HIRs preferentially occurred when animals were stationary, while HOR prevalence was less dependent on measured behaviors, consistent with previous reports^26^. While the theta oscillation is typically associated with running behavior, it has been suggested that theta oscillations can occur independent of movement, potentially underlying unique computation/brain states^4,44,45^. It is possible that HORs consist of multiple subgroups of HFOs, potentially dependent on distinct theta oscillations^4,46^.

The medial entorhinal cortex (MEC) supports routing of specific information to the hippocampus during theta oscillations, contributing to place cell dynamics and memory behaviors. It is thus likely that the interaction between gRSC and HPC during online states and HORs is also coordinated by MEC. HORs occur on the descending phase of HPC theta, accompanied by phased locked ‘nested’ gamma oscillations, consistent with spatial navigation and theta phase precession literature^4,47–49^. Gamma power was higher in the high- versus low-gamma band, likely driven by MEC inputs to CA1 Str. L.M.^4,33^, evidenced by a characteristic current sink in Str. L.M. locked to local CA1 theta oscillations (**Fig. 6B, right, black circle**). These results suggest the coordination of hippocampal high-gamma and HORS may be due to direct reciprocal connectivity between the gRSC and MEC^22,23,50,51^.

In conclusion we have demonstrated discrete HFO events occur in multiple behavioral states and are localized to L2/3. L5 can support HFO generation but spontaneous L5 HFOs do not occur due to a lack of sufficient excitatory inputs during CA1 theta and SWRs. Importantly, HORs are aligned with both gRSC and CA1 theta, strongly coupled to MEC-derived CA1 high gamma, suggesting a role for MEC in priming CA1-gRSC circuits for plasticity in following offline states. gRSC HFOs may be a general mechanism supporting communication and plasticity between hippocampus and cortex.

### Animal subjects

All data used in this study was collected in 12 sessions from 10 mice, aged 8-20 weeks. For optical stimulation of the gRSC layer experiments we used N = 3 C57BL/6J-CamKii-Cre:: C57BL/6J-Ai32 mice, for all other experiments N = 7 C57BL/6J:: FVB/NJ were used. Animals were housed in a 12-hour reverse light-dark cycle. All experiments were approved by the Virginia Tech IACUC.

## Methods

### Surgeries and habituation

Mice were anesthetized with isoflurane for all surgeries and received both local (bupivacaine 0.1 ml at 5 mg/kg) and systemic (meloxicam 0.1 ml at 5mg/kg) analgesia. In the first surgery a ground wire was implanted parallel with the surface of the cerebellum and a titanium head-bar was fixed above the skull and connected to the ground wire. The entire skull was then covered with dental cement (UNIFAST LC acrylic cement). Animals were monitored daily post-surgery. The day before recording craniotomies were made above the HPC and gRSC as follows (all measurements are made with respect to Bregma). HPC: 1.5mm M/L by −1.75 A/P; gRSC linear probe recordings: .5mm M/L on contralateral side of HPC craniotomy by −2.4 – −2.7mm A/P; gRSC opto-stimulation recordings: .5 – 1.25mm M/L by −2.4 – −2.7mm A/P. All animals were perfused within 24 hours of the end of the recording and 48 hours after perfusion brains were sectioned, mounted, and imaged. Animals were habituated to head fixation over the course of a 1-2 weeks then were water deprived to motivate them to run for water rewards in a 1-D virtual reality system. Mice learned this behavior in 7 days.

### gRSC and HPC recording setup

7 male and female, C57/B6J and C57BL/6J:: FVB/NJ mice were used in these experiments. Cambridge Neurotech’s H3 acute silicon electrodes were inserted into the craniotomies described above across the midline into the gRSC at 42-degree angle to the midline and a second silicon probe was implanted into ipsilateral HPC CA1. Probe positions were determined online using electro-anatomical landmarks and their position was confirmed post-hoc by imaging Di-I (applied to the Si probes before recording). For a subset of experiments electrolytic lesions were made using selected channels to further confirm the layer from which each electrode was placed during the experiment.

### gRSC Optostimulation recording setup

3 male and female, C57BL/6J-CamKii-Cre:: C57BL/6J-Ai32 mice were used in these experiments. All mice were head-fixed as described above, and Neurolight Technologies µLED probes were then implanted in the gRSC such that each shank was within each layer of the gRSC. To accomplish this the probe was set at a 15-degree angle from the midline. Electro-anatomical landmarks were used to identify recording depth during probe implant. Once the probe was in place optical stimulation intensity was empirically determined and set to an intensity that reliable evoked HFOs in L2/3. We then chose two lower intensities and gave 150, 100ms pulses of each low, medium, and high intensity stimulations each with a randomized delay of 1-3 seconds between stimulations (only high intensity stimulations shown in text; **Fig. 2**). Di-I fluorescence was used to post hoc identify probe implant locations.

### Tissue processing and immunohistochemistry

Within 24 hours after recordings, animals were perfused using 4% paraformaldehyde. Brains were extracted and sliced 48-hours after perfusion. Sections were mounted with DAPI containing mounting solution and imaged using a confocal microscope.

### Data acquisition

Silicon probes from Cambridge Neurotech*, IDAX*, or Neurolight Technologies** were used to obtain LFP and single unit recordings. Electrophysiology data were acquired using an Intan RHD2000 recording system (sampled at 30 kHz).

* = gRSC and HPC dual recordings

** = opto-stimulation recordings

## Data analysis

All recordings were down sampled to 1.25kHz to analyze LFP activity. Statistical analyses were conducted using MATLAB (Statistical Toolbox, Circular Statistics Toolbox, CellExplorer, and Buzcode Toolbox). Sample sizes were not determined by any statistical analyses but are consistent with sample sizes used in previously published studies in the field. Data is shown as mean ± s.e.m. with significance determined by using Wilcoxon Signed Rank test unless otherwise stated in the text.

### HFO and SWR event detection

For high frequency oscillation detection, channels from the gRSC were chosen with the highest power in the ~150Hz range across the duration of the entire recording. For sharp wave-ripple detection, channels from the HPC with the highest power in the ~150 Hz range were chosen by visual inspection. Selected channels were bandpass filtered between 100-250 Hz. HFO events were identified as events no longer than 150 ms that surpassed 1 SD above the mean of the rectified and filtered signal. SWR events were identified as events no longer than 150 ms that surpassed 3 SD above the mean of the rectified and filtered signal. Each group of events detected from each recording were visually inspected using Neuroscope2. HFO events were then split into HIRs and HORs. HIRs are HFOs that occur within +/− 50 ms of an HPC SWR, and HORs were all other events outside +/−50ms of SWRs (**Fig. 3**).

### Unit classification, monosynaptic connection identification, and HFO response

Unit activity was extracted from the raw 30 kHz recordings using Kilosort1 and manually curated in Phy2. Units were putatively labelled excitatory or inhibitory units using CellExplorer^31^. Putative monosynaptic connections were also determined in CellExplorer using established methods and manually checked for accuracy^52–54^. Gain in single unit firing rate to HORs and HIRs was calculated as the average in-event firing rate divided by the baseline firing rate outside of all events. Percentages of cells making monosynaptic connections were calculated by finding the number of ENs (E-E or E-I) and INs (I-E or I-I) making intra-laminar connections. The number of units making these connections was then divided by the total number of that presynaptic unit type found in the.

### Spectral analysis

For the averaged and aligned whole recording power spectral density (PSD) (**Fig. 1**) all channels were whitened, and low pass filtered at 250 Hz. A fast Fourier transform was then computed and smoothed using a Savitzky-Golay filter for each channel. For event triggered PSDs (**Figs. 5 & 6**) channels used to detect HFOs and SWRs were selected and whitened. A sliding 16 ms window with 8ms steps was for each event. At each window step a Morlet Wavelet decomposition was then performed for integer frequencies of 1-500 Hz.

### Current Source Density analysis

Current source density plots (**Figs. 5 & 6**) were created by finding the second spatial derivative over all electrodes on the probe at each time point around the events. All HIR or HOR triggered CSDs were averaged together for both gRSC and HPC recordings.

### HOR theta phase preference

For each session the channel used to detect HFOs and SWRs were selected, bandpass filtered from 6-10 Hz, and a Hilbert transform was performed on each. The phase of the signal at each time point was then calculated and theta phase at the peak of each HOR was found. The Rayleigh’s test was used to determine significance (**Fig. 5**).

### Power-Power comodulation analysis

Channels used to detect HFOs and SWRs were selected and a fast Fourier transform was performed in sliding 2 second windows with 1 second steps. The Spearman correlation for each frequency bin from the gRSC was found for the comparison to every frequency bin in the HPC over the duration of the entire recording (**Fig. 6**).

## Contributions

K.C.A., D.F.E. and S.M. designed the study. K.C.A. conducted all aspects of the study. E.T.G., L.M.F.K. J.K. C.M.B. and J.C.B. assisted with data analyses, and interpretation. All the authors contributed to writing the manuscript.

## ACKNOWLEDGMENTS

D.F.E. is supported by grants from The Simons Foundation and The Whitehall Foundation. S.A.M. is supported by NIMH (R00MH11842). K.C.A. is supported by the Virginia Tech School of Neuroscience and the Virginia Tech Molecular and Cellular Biology program.

