## Supplementary Figs for "Granular retrosplenial cortex layer 2/3 generates high-frequency oscillations coupled with hippocampal theta and gamma in online states or sharp-wave ripples in offline states"

### Supplementary Figures

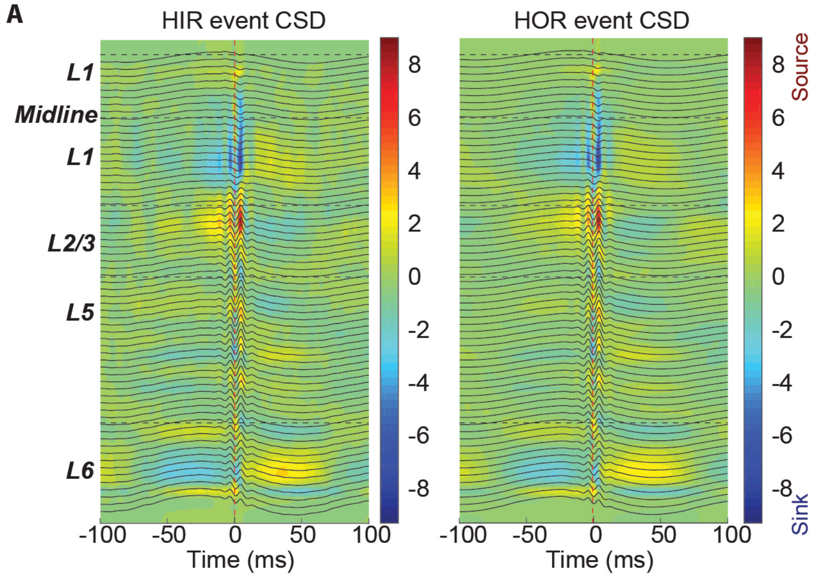


**Supplementary Fig. 1: HIRs and HORs detected in gRSC layer 5: A)** Current source density analysis of HIRs and HORs detected in layer 5 of the gRSC. Note current sources and sinks are still located at the L1 and L2/3 border.

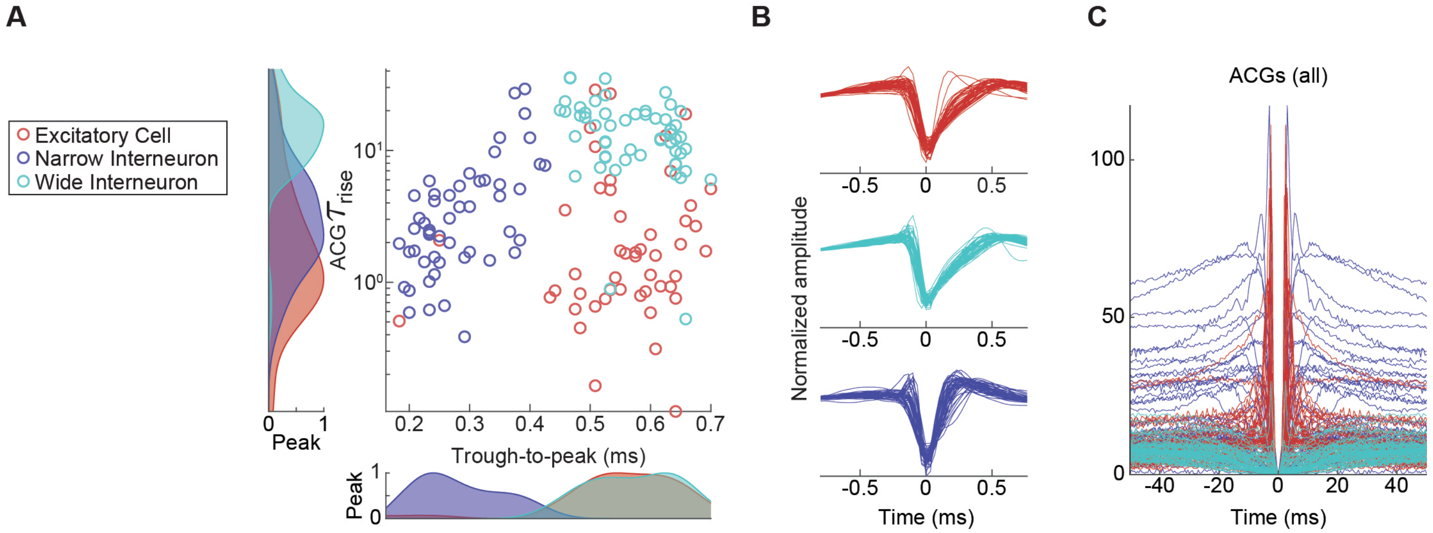


**Supplementary Fig. 2: Putative cell type classification of gRSC neurons: A)** Scatter plot of unit auto-correlogram (ACG) tau rise and wave form trough-to-peak duration, used to classify cells as excitatory (red), narrow interneurons (dark blue), or wide interneurons (light blue). **B)** Average wave forms of each cell, split by type. **C)** ACGs of each cell color coded to match cell type as in A.

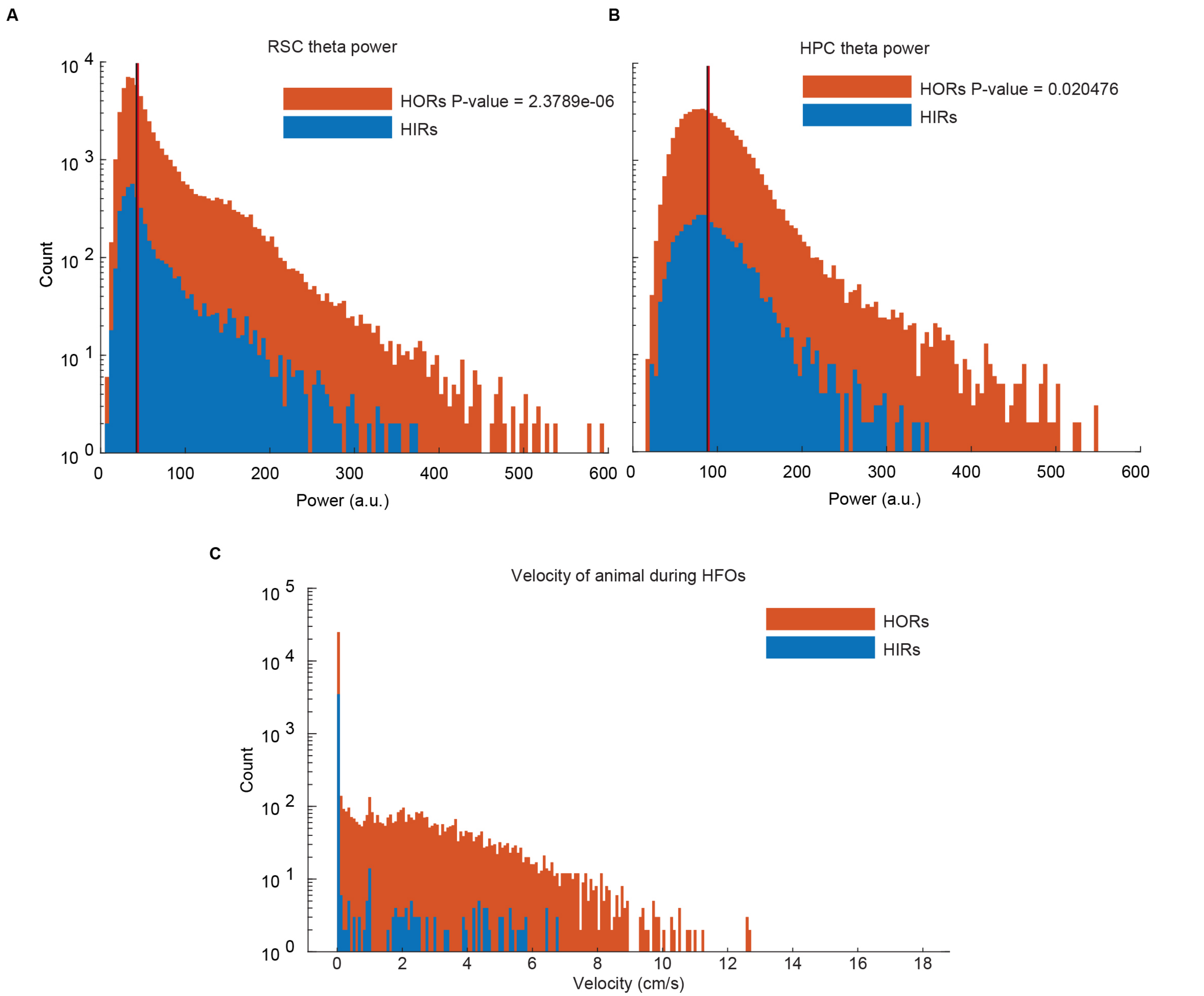


**Supplementary Fig. 3: Theta power and velocity during HIRs and HORs: A)** gRSC theta power during HFO events, HORs (red line = median theta power during HORs) have a significantly higher theta power than HIRs (red line = median theta power during HIRs). **B)** Same as A but for HPC theta power, HORs have a significantly higher theta power than HIRs. **C)** Velocity of the animal during HFO events. **Fig. 5C** shows the percent of events occurring during movement and no movement.
